# HiFi-SIM: reconstructing high-fidelity structured illumination microscope images

**DOI:** 10.1101/2020.07.16.204875

**Authors:** Gang Wen, Simin Li, Linbo Wang, Xiaohu Chen, Zhenglong Sun, Yong Liang, Xin Jin, Yuguo Tang, Hui Li

## Abstract

Structured illumination microscopy (SIM) has been a widely-used super-resolution (SR) fluorescence microscopy technique, but artifacts often appear in reconstructed SR images which reduce its fidelity and might cause misinterpretation of biological structures. We present HiFi-SIM, a high-fidelity SIM reconstruction algorithm, by engineering the effective point spread function (PSF) into an ideal form. HiFi-SIM can effectively reduce commonly-seen artifacts without loss of fine structures and improve the axial sectioning. Since results of HiFi-SIM are not sensitive to used PSF and reconstruction parameters, it lowers the requirements for dedicated PSF calibration and complicated parameter adjustment, thus promoting SIM as a daily imaging tool.

---

Despite the relatively modest improvement in resolution compared to that in other single-molecule-localization or emission-depletion-based SR techniques, SIM are recognized as a most promising tool for studying the dynamics of subcellular structures in live cells due to its high photon-efficiency, little photon damage and bleaching, as well as compatibility with most fluorescent labeling protocols^1–5^. However, since the final SR images rely heavily on post-processing algorithms that are prone to reconstruction artifacts, the fidelity and quantification of SR-SIM are always challenged^4–7^. Many studies have been conducted aiming at reducing the artifacts in SIM images, by instrument refinement and proper maintenance^8^, establishing standard experimental protocol^6^, and developing different image reconstruction algorithms^9–14^ Despite all the efforts, artifacts still remain^6,7^ hence limiting the application of SIM as a daily imaging tool. Further, new structures discovered using SR-SIM need to be interpreted with special care to avoid misinterpretation of artifacts as real features.

By now the reconstruction algorithms used by most commercial SIM setups and open source packages such as Wiener-SIM in SIMToolbox^15^, FairSIM^16^, and OpenSIM^17^ were based on principle of the Wiener-SIM algorithm established by Gustafsson^10^. Reconstruction of SR-SIM is conducted by recombination of different spectrum components in frequency domain, which inevitably leads to an effective optical transfer function (OTF) with non-smooth shape (**Fig. 1a, c, e**). By inverse Fourier transformation, the downward kinks in the OTF are transformed into sidelobes in the PSF (**Fig. 1b, d, f**). The abnormal features of OTF could be corrected by the generally used Wiener deconvolution procedure, but that requires the used OTF and estimated reconstruction parameters accurately match the actual imaging conditions^1,6^. Otherwise, the final SR images will exhibit obvious reconstruction artifacts. Further, the raised peaks in OTF enhance the residual out-of-focus signal, hence limit the optical sectioning capability and cause honeycomb artifacts^2,3^. Consequently, even for high-quality data acquired using well-maintained setup and appropriate samples, artifacts cannot be totally avoided in SIM images, indicating the inherent defects of Wiener-SIM (**Supplementary Note 1**).

**Fig. 1.**
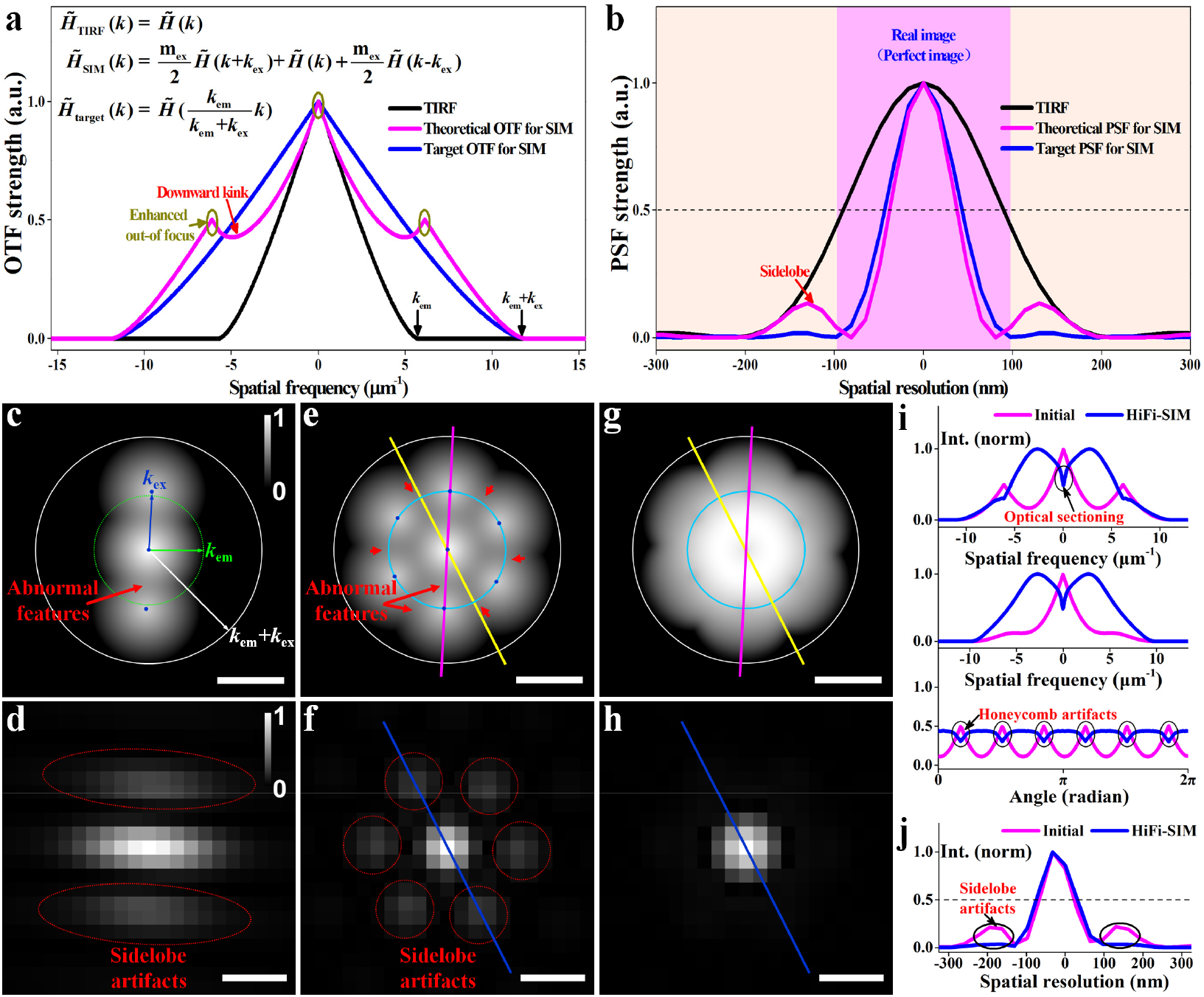
Principle of OTF optimization in HiFi-SIM. **a,b,** Theoretical OTFs and corresponding PSFs for wide-field fluorescence microscopy (total internal reflection fluorescence microscopy, TIRF) and directly-combined SR-SIM, and the ideal target OTF and corresponding PSF for SR-SIM. The downward kinks in the OTF of directly-combined SR-SIM result in sidelobes in PSF, while the upward peaks enhance out-of-focus signals which eventually limit the optical sectioning and cause honeycomb artifacts. **c,d**, 2D OTF and corresponding PSF of directly-combined SR-SIM with excitation pattern in only one orientation. **e,f**, 2D OTF and corresponding PSF of directly-combined SR-SIM with excitation patterns in three orientations. Green and white circles represent the diffraction limited boundaries of the wide-field and SR-SIM, respectively; blue spots represent the spatial frequencies of the excitation pattern at different orientations; cyan circles represent the circular cross-section with a radius equal to the spatial frequency of the excitation patterns. **g,h,** Equivalent OTF and corresponding PSF after optimizing the directly-combined OTF by HiFi-SIM. **i**, Intensity profiles along the magenta, yellow, and cyan lines in **e** and **g. j**, Intensity profiles along the blue lines in **f** and **h**. Gamma value: 0.3 for (**c**), (**e**) and (**g**). Scale bar in **c,e,g** 7 μm^−1^; in **d,f,h,** 150 nm.

The ideal form of OTF for SR-SIM should be same form of OTF for wide-field imaging 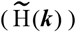 with the cut-off frequency extended to (***k***_em_ **+ *k***_ex_), where ***k***_em_ is the cut-off frequency of wide-field imaging and ***k***_ex_ is spatial frequency of the excitation pattern (**Fig. 1a**), so that the reconstruction artifacts could be diminished in principle. We present a high-fidelity SIM reconstruction algorithm, called “HiFi-SIM”, by engineering the effective OTF to the target form. The reconstructed spectrum obtained by directly recombining different spectrum components is first optimized by an optimization function *W*_1_(***k***) with the following expression

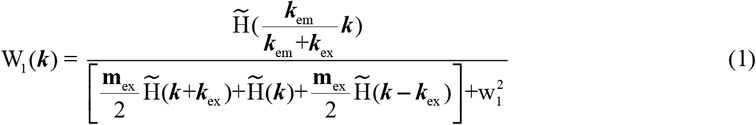

where w_1_ is an empirical constant. Then, a deconvolution optimization function W_2_(***k***) is further applied. In the above two steps, an OTF attenuation is applied as the weight function to balance the tradeoff between suppressing out-of-focus signal and preserving the fine structures,^11,16^. With the spectrum optimization, an equivalent OTF with uniform axial and circumferential distribution can be obtained, thereby realizing perfect SR-SIM with little artifacts by HiFi-SIM (**Fig. 1g, h, i, j**).

By combining the normalized cross-correlation method^4^ with a spectrum notch, HiFi-SIM could automatically estimate the pattern parameters from most raw data, including low SNR data, TIRF-SIM data, and even data with obvious periodic structures (**Supplementary Note 2**). Moreover, HiFi-SIM is not sensitive to used OTF and reconstruction parameters^6, 18^ (**Supplementary Note 3**). For most tested data with acceptable quality, only the OTF attenuation strength parameter needed to be adjusted during reconstruction (**Supplementary Note 4**). Thus, HiFi-SIM could produce high-fidelity SR images with no observable artifacts, requiring minimal user intervention, hence promoting wide application of SIM as a daily imaging tool.

To quantitatively evaluate the fidelity of HiFi-SIM, three representative structures (spherical beads, lines, and rings) were imaged by a custom-built laser-interference SIM^19^ and GE DeltaVision OMX. SR images were reconstructed with the traditional Wiener-SIM, open source package fairSIM, GE SoftWoRx, and HiFi-SIM (**Fig. 2**). The reconstructed spectrum by traditional Wiener-SIM and fairSIM displayed patchy features; therefore, symmetrically distributed snowflake-like artifacts appeared around the real beads in the SR images (**Fig. 2a, b**). Artificial lines with intensity approximately 10%–30% of the actual structures also appeared as sidelobe in the SR images obtained by SoftWoRx and fairSIM (**Fig. 2h, i, l, o, p, s**). In contrast, HiFi-SIM corrected the abnormal spectrum features, yielding clean bead, and line and ring structures (**Fig. 2c, j, q**). The root mean square error (r.m.s.e.) of the error-map, between the reconstructed image using HiFi-SIM and the ground-truth model, was decreased approximately 4 times (**Fig. 2m**). The calculated structural similarity remained above 91.3% within different reconstruction parameters of HiFi-SIM, indicating the reconstruction parameters to not require routine adjustment (**Fig. 2t**).

**Fig. 2.**
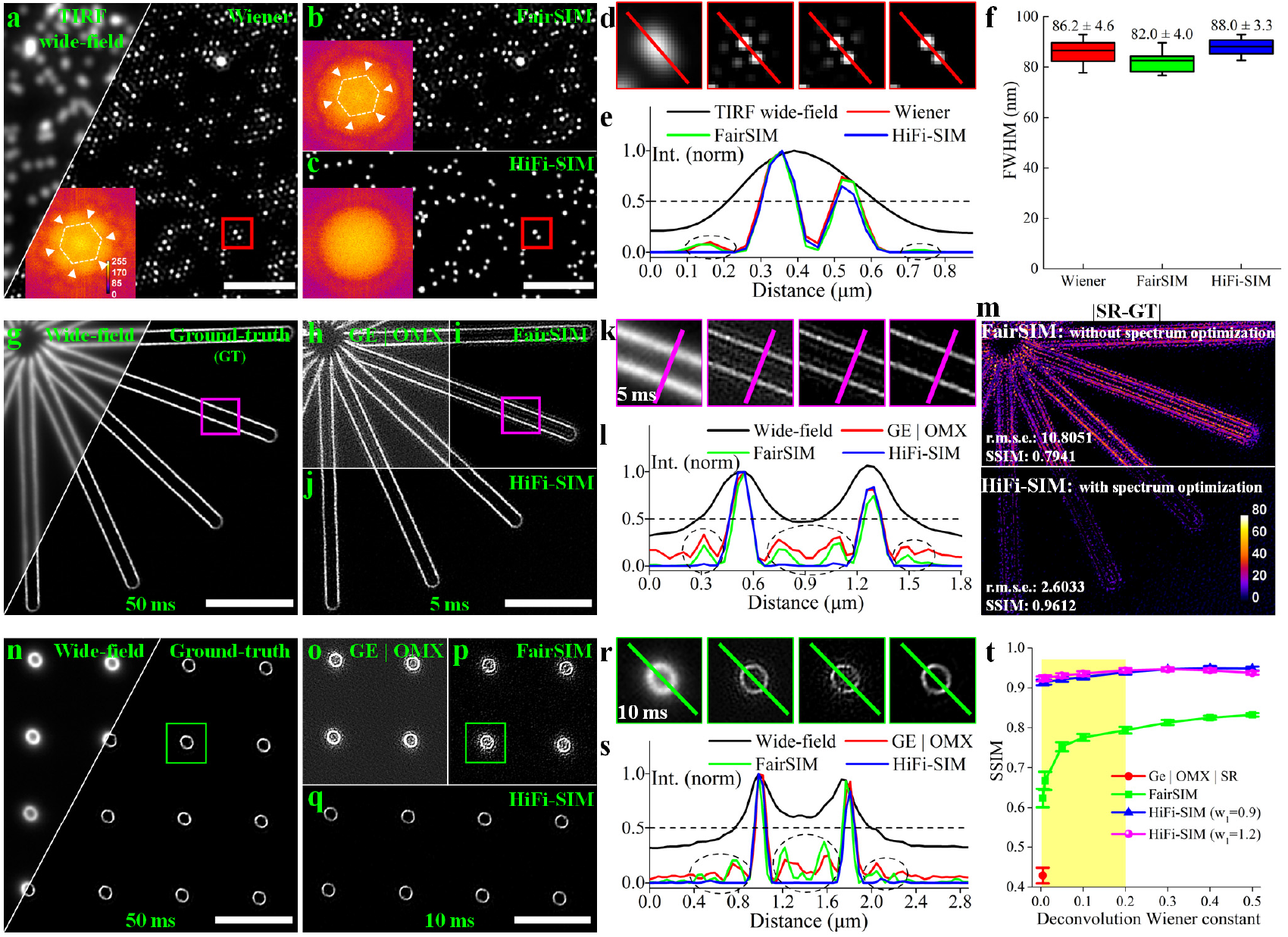
Quantitative evaluation of the fidelity of HiFi-SIM reconstruction. **a-c,** SR images of fluorescent beads reconstructed by the traditional Wiener-SIM, fairSIM, and HiFi-SIM; insets show the corresponding reconstructed spectrum; **d**, Magnified images of the red-box regions in **a-c. e,** Line profiles along the beads show diminishing sidelobe artifacts in HiFi-SIM. **f**, Full-width half-maxima (FWHMs) of the fluorescence profiles of 10 beads in **a-c. g**, Wide-field and ground-truth (GT) images of “start” patterns in Argo-SIM slide, seen as typical line structures (see **Methods**). **h-j,** SR images of “start” patterns with 5-ms exposure time reconstructed by SoftWoRx, fairSIM, and HiFi-SIM. **k,** Magnified SR images of “start” patterns reconstructed from the magenta-box regions. **l,** Intensity profiles along the magenta lines in **k. m**, Error-map of the SIM images to the ground-truth model. Root mean square error (r.m.s.e.) and structural similarity index (SSIM) values are shown in the left-lower corner. **n-s,** Comparison of the SR images for ring patterns in Argolight slide, reconstructed by SoftWoRx, fairSIM, and HiFi-SIM. **t**, SSIM plot of reconstructed SR results vs. the ground-truth model, with different reconstruction parameters. Scale bar in **a-c,** 2 μm; in **g-i** and **m-p,** 5 μm.

We further imaged microtubules in live COS-7 cells using TIRF-SIM mode, and the raw data were reconstructed with fairSIM and HiFi-SIM (**Fig. 3a-g**). High modulation contrast enabled both algorithms to reconstruct good SR image. However, with fairSIM, sidelobe artifacts with intensity approximately 10% of that in actual microtubules were present in certain regions, and overlap of two sidelobes from nearby microtubules created a structure with double intensity, which could very likely be interpreted as a real structure (arrows in **Fig. 3e**). With HiFi-SIM, the abnormal spectrum could be effectively corrected to eliminate artifacts, thereby leaving no observable sidelobe (**Fig. 3c, f, g**). Similarly, SR image of microtubules in fixed COS-7 cells, reconstructed by SoftWoRx showed significant residual out-of-focus background and artifacts, whereas HiFi-SIM could effectively remove the background fluorescence and reduce artifacts (**Supplementary Fig. 1**).

**Fig. 3.**
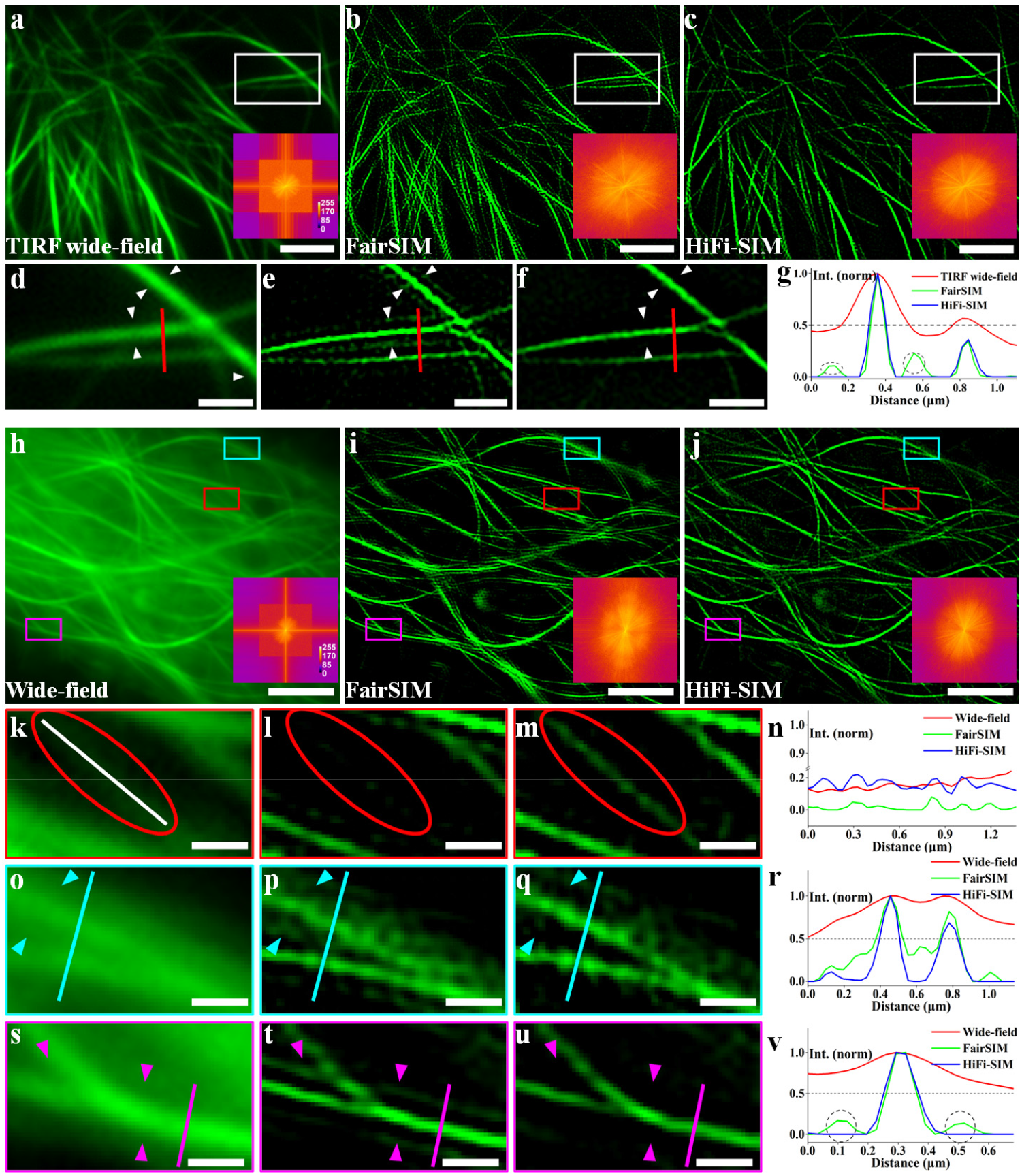
Performance of HiFi-SIM on reconstructing raw data with high-quality (a-g) and with suboptimal quality (h-v). **a**, Wide-field image of microtubules in live COS-7 cells. **b, c**, SR-SIM images, reconstructed by fairSIM and HiFi-SIM, for raw data collected under well-aligned TIRF-SIM condition. Right-lower corner shows corresponding spectrum. **d-f**, Magnified SR images of the white-box regions in **a-c. g,** Intensity profiles along the red lines in **d-f. h**, Wide-field image of microtubules in live U2OS cells with strong background fluorescence. **i, j**, SR images reconstructed by fairSIM and HiFi-SIM for raw data with low-modulation depths in three orientations. **k-m**, Magnified images of the red-box regions in **h-j. n**, Intensity profiles along the white lines in **k-m. o-q**, Magnified images of the cyan-box regions in **h-j. r**, Intensity profiles along the cyan lines in **o-q. s-u**, Magnified images of the magenta-box regions in **h-j. v**, Intensity profiles along the magenta lines in **s-u**. Scale bar in **a-c**, 3 μm; in **d-f**, 1 μm; in **h-j**, 4 μm; and in **k-u**, 0.5 μm.

In Wiener-SIM, increasing the OTF attenuation strength or Wiener constant can reduce certain artifacts^6,16^; however, it may cause loss of fine or weak structures (**Supplementary Note 1**). This situation would get worse for low-modulation data. To mimic sub-optimal SIM imaging, microtubules in live U2OS cells were imaged (**Fig. 3h-v**). The raw data were collected in SIM mode with incident beam angle smaller than the critical angle of TIRF, and the modulation depths were intentionally adjusted to be different by controlling the beam polarization^8^. In practice, such suboptimal situations often occur in SIM imaging for different kinds of reason and the data were generally abandoned These data with low modulation depths (0.43, 0.36, and 0.21 in three orientations) and strong background are suggested to be reconstructed with special care by fairSIM, since serious honeycomb and sidelobe artifacts as well as loss of tiny structures are inevitable (**Fig. 3i, l, p, t**). In comparison, HiFi-SIM reconstructed SR image with little honeycomb and sidelobe artifacts, and the lost tiny structures were preserved (**Fig. 3j, m, q, u**). Hence, HiFi-SIM could better balance the trade-off between suppressing artifacts and preserving sample structures, such that many previously abandoned suboptimal data could be reused.

Out-of-focus background is another challenge that limits the reconstruction quality of two-dimensional (2D) SIM^2, 3, 5^. HiFi-SIM was also applied to reconstruct volume stack of 2D-SIM data collected on DeltaVision OMX. SR images of microfilaments in fixed U2OS cells by HiFi-SIM showed higher contrast and fewer artifacts, indicating its better sectioning capability compared to SoftWoRx (**Supplementary Fig. 2**). HiFi-SIM has also been extended to reconstruct single-layer 3D-SIM datasets (**Supplementary Note 5**), and its reconstruction quality was comparable to the quality of the same layer in full 3D-SIM reconstruction (**Supplementary Fig. 3**).

We extensively tested HiFi-SIM with different datasets, including caveolae in live cells (**Supplementary Fig. 4**), hollow vesicles in live cells (**Supplementary Fig. 5**), endoplasmic reticulum with strong background (**Supplementary Fig. 6**), and dataset from open sources (**Supplementary Figs. 7-9**). With several other available algorithms as benchmarks, HiFi-SIM was demonstrated to yield better quality of SR images in all tested cases.

In conclusion, HiFi-SIM provides an easy-to-use SIM reconstruction package yielding high-fidelity SR images from commercial and home-built SIM setups, with good-to-bad quality data. HiFi-SIM can be easily extended to multi-color imaging, to combine with rolling-SIM^4^, or to conduct GPU acceleration^20^. The spectrum optimization principles in HiFi-SIM can also be further applied to reconstruct images from non-linear SIM^2^ or lattice light-sheet microscopy^21^, which are more prone to unnatural spectrum and reconstruction artifacts.

## Methods

Methods, including statements of data availability and any associated accession codes and additional references, are available in the online version of the paper.

## Acknowledgments

We thank Prof. Dr. Thomas Huser and Dr. Marcel Müller (Bielefeld University) for the fruitful discussion on the algorithm. The work was supported by the National Key Research and Development Program of China [grant no. 2017YFC0110100] and the National Natural Science Foundation of China [grant no. 11504409].

## Author contributions

G.W. and S.L. initiated the spectrum optimization idea; G.W. and L.W. developed the reconstruction algorithm; G.W. performed the experiments and analyzed the data; S.L., X.C, Y.L., and X.J. built the SIM imaging system; Z.S. performed sample preparation; H.L. and Y.T. conceived and supervised the study; G.W. and H.L. wrote the paper. All the authors participated in the discussions and data interpretation.

## Methods

### Standard fluorescent sample

Fluorescent beads of 100-nm diameter and commercial Argo-SIM slide were employed as standard samples to quantitatively evaluate the fidelity of reconstruction algorithms. Carboxylate-modified beads (0.1 μm, yellow-green fluorescent 505/515, F8803) were purchased from Thermo Fisher Scientific (MA, USA) and diluted 100 times before use. Commercially available coverslips (~150 μm thick) with 24 × 60 mm (Cellvis, USA) were carefully cleaned using the procedure in [20]. A silicone sheet (GBL665201-25EA, Sigma-Aldrich, USA) with a well of 9-mm diameter was attached to the coverslip. The fluorescence solution was dispensed onto the coverslip and imaged in PBS buffer. The “Star” and “2D matrix of rings” patterns in Argo-SIM were used as typical “line” and “ring” structures.

### Cell culture and labeling

COS-7 and U2OS cells were obtained from the Cell Bank of the Chinese Academy of Sciences (Shanghai, China) and cultured in an incubator at 37 °C and 5% CO_2_. The COS-7 cells were cultivated in a DMEM medium (Thermo Fisher Scientific, USA) supplemented with 1% penicillin G, streptomycin (Sangon Biotech, China), and 10% fetal bovine serum (Thermo Fisher Scientific, USA). The U2OS cells were cultivated in McCoy’s 5A medium, modified (Thermo Fisher Scientific, USA), supplemented with 1% penicillin G, streptomycin (Sangon Biotech, China), and 10% fetal bovine serum (Thermo Fisher Scientific, USA).

Cells were transiently transfected using Lipofectamine 2000 (Thermo Fisher Scientific, USA) as per manufacturer’s protocol. The mEmerald-Tubulin-N-18 vector (plasmid #54293, Addgene, USA), mEGFP-Lifeact vector (plasmid #54610, Addgene, USA), mEmerald-Caveolin vector (plasmid #54025, Addgene, USA), and mEmerald-ER-3 vector (plasmid #54082, Addgene, USA) were used to label the microtubule, microfilament, caveolae, and endoplasmic reticulum, respectively. Cell vesicles were labeled by the CD63-EGFP vector, which was constructed by inserting homo sapiens CD63 cDNAs into pEGFP-n1 vector (Clontech, USA). Twenty-four hours after transfection, the cells were detached using trypsin-EDTA (Thermo Fisher Scientific, USA), seeded onto poly-L-lysine-coated 35-mm glass-bottom dishes (Cellvis, USA), and cultured in an incubator at 37 °C and 5% CO_2_ for an additional 24 h before the experiments.

For live cell imaging, the complete medium was replaced by HBSS solution (Thermo Fisher Scientific, USA) containing Ca^2+^ and Mg^2+^ but no phenol red. For fixed cell imaging, the complete medium was removed and cells were fixed with 4% paraformaldehyde for 10 min at room temperature. After fixation, cells were washed thrice by PBS buffer. Both live and fixed cells were imaged in PBS buffer.

### SIM imaging

SIM experiments were performed using commercial SIM microscopes, namely DeltaVision OMX SR (GE Healthcare in Issaquah, Washington, USA) and N-SIM S (Nikon Corporation, Tokyo, Japan), as well as a custom-built two-beam interference SIM microscope (**Supplementary Fig. 10**). The custom-built SIM was constructed around a commercial inverted fluorescence microscope (IX83, Olympus Life Science, Japan) with a TIRF-oil-immersion objective (UAPON 100×, NA = 1.49, Olympus Life Science, Japan). A 488-nm, 500-mW semiconductor laser (Genesis MX488-500 STM, Coherent, USA) was used for excitation, a quad-band total internal reflection (TIRF) filter block (TRF89902-EM, Chroma, USA) was employed for imaging, and a sCMOS camera (ORCA-Flash 4.0 V2, Hamamatsu, Japan) was used as the detector. To generate illumination patterns with different periods, a ferroelectric liquid-crystal spatial light modulator (SLM, SXGA-3DM, Fourth Dimension Displays, UK) was employed as the grating.

Beads with 100-nm diameter, microtubules, and microfilaments in live COS-7 cells were imaged using the custom-built setup in TIRF-SIM mode. Microtubules and vesicles in live U2OS cells, and endoplasmic reticulum in fixed U2OS cells were imaged in the setup under conventional SIM mode, with incident beam angle smaller than the critical angle of TIRF. Reconstruction parameters: for beads data, emission wavelength (λ_em_) = 515 nm while that for microtubule, microfilaments, vesicles, and endoplasmic reticulum data was 525 nm, and the single pixel size of the detector is calibrated to 65 nm/pixel.

The “**2D matrix of rings**” and “**Star**” patterns in Argo-SIM, microtubules in fixed COS-7 cells, and microfilaments in fixed U2OS cells were imaged on the DeltaVision OMX SR with the parameters: NA=1.42 (oil immersed), excitation wavelength (λ_ex_) = 488 nm, emission wavelength (λ_em_) = 527 nm, and the single pixel size of the detector is calibrated to 78.6 nm/pixel. In addition, caveolae in live U2OS cells were imaged on the N-SIM S with the parameters: NA=1.49 (oil immersed), excitation wavelength (λ_ex_) = 488 nm, emission wavelength (λ_em_) = 525 nm, and the single pixel size of the detector is calibrated to 60 nm/pixel.

### SIM reconstruction with commercial and open-source software

Commercial SIM reconstruction software packages, including GE SoftWoRx and Nikon NIS-Elements, and open source packages, including fairSIM, Hessian denoising SIM, Total Variation (TV) denoising SIM, and Maximum a posteriori probability SIM (MAP-SIM) in SIMToolbox were used for comparative super-resolution image reconstruction. Images labeled ‘GE | OMX’ were reconstructed with SoftWoRx, and the Wiener constants were 0.005 by default. Images labeled ‘Nikon | N-SIM | 2D’ were reconstructed with NIS-Elements. Images labeled ‘FairSIM’ and ‘RL-SIM’ were reconstructed with the Wiener-SIM and RL-SIM in fairSIM, respectively. The adopted OTFs were approximate OTFs (“dampening” factor = 0.3), and Wiener constants defaulted to 0.1. Images labeled ‘SIM | Hessian denoised’ were obtained by additional Hessian denoising of the reconstructed results of fairSIM. Images labeled ‘SIM | TV denoised’ were obtained by additional TV denoising of the reconstructed results of fairSIM. Images labeled ‘Map-SIM’ were reconstructed with the Map-SIM in SIMToolbox. The test dataset was downloaded from the official website of SIMToolbox, and the same reconstruction parameters, as in the paper, were set. As a comparison with HiFi-SIM, we have also implemented a traditional Wiener-SIM. In the implementation, raw data preprocessing, and reconstruction parameter estimation use the same method as in HiFi-SIM, while the recombination of spectrum components follows the traditional Wiener deconvolution procedure (**Supplementary Note 1**). Images labeled ‘Wiener’ were reconstructed with the traditional Wiener-SIM we implemented.

### SIM reconstruction with HiFi-SIM

The flow chart of HiFi-SIM algorithm is shown in **Supplementary Fig. 11a**, which includes: (1) raw data preprocessing, (2) automatic parameter estimation, and (3) spectrum recombination and optimization. The details are described below.

#### (1) Raw data preprocessing

To avoid complicated calibration of the system OTF during SIM imaging, which is not practical for most ordinary SIM users, HiFi-SIM uses an approximate OTF generated based on the objective used and emission wavelength^18^

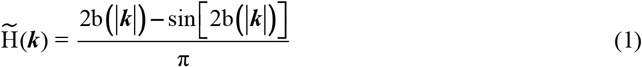

where *b***(*k*)**=cos^−1^ **(*k/k***_em_), and ***k***_em_ = 2NA/**λ**_em_ is the cutoff frequency of the imaging system in reciprocal space; NA is numerical aperture of the detection objective; **λ**_em_ is the emission wavelength.

The PSF used for raw SIM data preprocessing in HiFi-SIM was obtained by Fourier transform of the above OTF.

To enhance the peaks of pattern wave vectors in the cross-correlation map and remove certain out-of-focus background and noise^12,22^, Richardson-Lucy deconvolution was first performed on raw data using the generated PSF in HiFi-SIM. The default iteration number was set as 5.

#### (2) Automatic reconstruction parameter estimation

Determining the correct reconstruction parameters from the raw data is essential for reconstructing the SR-SIM images with minimal artifacts. HiFi-SIM basically follows the “cross-correlation-based” method, implemented in fairSIM, to determine reconstruction parameters^16^. However, to enable automatic parameter estimation, without hand-defined regions in the cross-correlation map, HiFi-SIM adopts a method of combining amplitude normalized cross-correlation and spectrum notch to determine the pattern wave vectors (**Supplementary Fig. 11a**). Compared to the conventional cross-correlation method in fairSIM, and the normalized phase cross-correlation method in Hessian-SIM^4^, HiFi-SIM could automatically estimate the reconstruction parameters of most of the raw data precisely, even for that with obvious periodic structures (**Supplementary Fig. 12**).

#### (3) Spectrum recombination and optimization

HiFi-SIM first calculated the weighted sum of all the shifted spectrum components 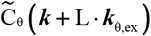 to obtain the directly-combined spectrum as follows

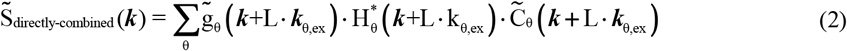

where

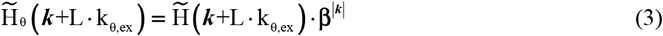

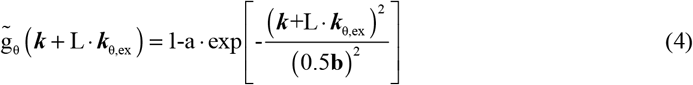

and θ and L represent the pattern orientation and shifted spectrum order, respectively. ***k***_θ,ex_ represents the excitation-pattern wave vector. **β** (0~1) is the “dampening” factor for compensating aberrations^8,16^. 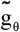 is a Gaussian function used to perform the OTF attenuation; **a** and **b** correspond to attenuation strength and width.

According to the frequency domain Wiener filtering^25^, the mathematical formula of optimization function W_1_(***k***) for initial optimization is as follows

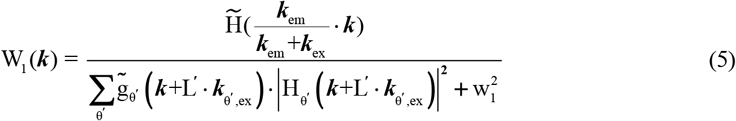

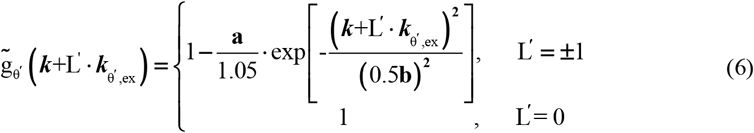

where w_1_, is the Wiener constant of the initial optimization, which is an empirical value. Here, the power spectrum 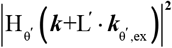 instead of 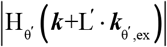 isused in the denominator to avoid errors arising from coordinates of **±k**_θ’,ex_, not satisfying the “Fourier shift theorem”^10, 17^. Specifically, W_1_(***k***) can not only correct the abnormal features of the directly-combined OTF, but also suppress the residual out-of-focus signal (**Supplementary Fig. 13b, e, h**).

The reconstruction result obtained by optimizing the directly-combined spectrum using W_1_ (***k***) is

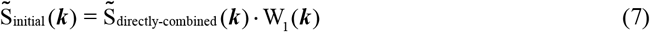

Further, deconvolution optimization could be performed with a function W_2_(***k***) as follows

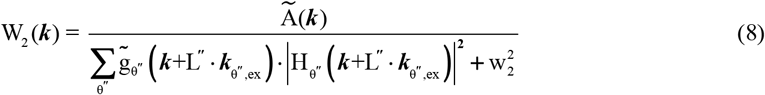

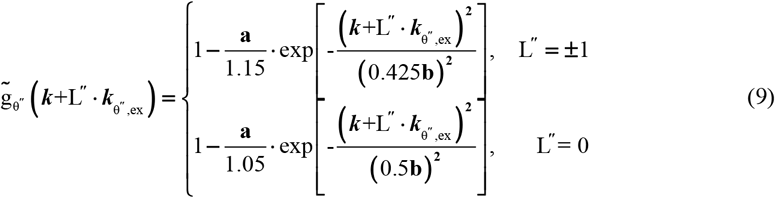

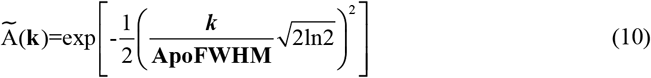

where w_2_ is the deconvolution Wiener constant and 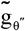 is a weighting function. **ApoFWHM** is the full width at half maximum (FWHM) of the apodization function. In particular, W_2_(***k***) was used not only for deconvolution, but also for recovering the real sample signals suppressed by OTF attenuation (**Supplementary Fig. 13c, f, i**).

The final high-fidelity SR image could be obtained by

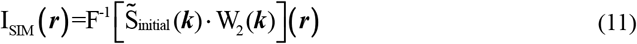

where symbol F^−1^ represents inverse Fourier transform. See **Supplementary Note 3** for a more detailed discussion of the spectrum optimization of HiFi-SIM.

Importantly, in HiFi-SIM, the default values of empirical parameters worked well for most of the data tested: **β**=1.0; **b**=1.0; **ApoFWHM**=min**{**0.5, 0.5***k***_ex_/***k***_em_**}**; w_1_=1.2; w_2_=0.1. Only attenuation strength **a** needed to be manually set to adjust the optical sectioning. For a detailed discussion of the reconstruction parameter settings, see **Supplementary Note 4**.

HiFi-SIM was implemented with a graphical user interface (GUI) of MATLAB software (**Supplementary software**).

### Simulation of OTF of SR-SIM

The OTFs and PSFs of wide-field imaging, directly-combined SR-SIM and ideal target SR-SIM (**Fig. 1a,b**) were simulated from the theoretical OTF of TIRF imaging and TIRF-SIM imaging condition (Eq. (**1**)). The imaging system parameters includes excitation wavelength of 488 nm, emission wavelength of 525 nm, oil objective with NA of 1.49, and the single pixel size of 65 nm/pixel.

The theoretical OTF of directly-combined SR-SIM was deduced as^2^

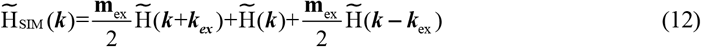

where **m**_ex_=1.0 is the modulation factor of the excitation patterns.

The ideal target OTF of SR-SIM has the same form as wide-field imaging but the cut-off frequency extended to ***k***_em_ **+ *k***_ex_, as follow

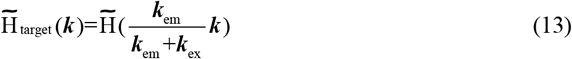

In 2D-SIM, the OTF corresponding to the directly-combined spectrum 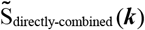 was generated based on the power spectrum of theoretical OTF (**Fig. 1c,e**)

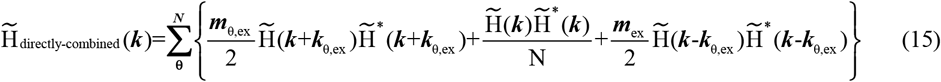

where θ represents the pattern orientation, N represents the total number of orientations.

The equivalent OTF of HiFi-SIM was obtained by optimizing 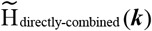 using W_1_(***k***) and W_2_(***k***) (**Fig. 1g**)

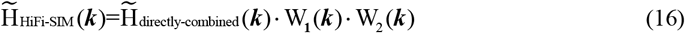

### Quantification of fidelity of SR images

To quantitatively evaluate the fidelity of SR images reconstructed by HiFi-SIM, two typical patterns (rings and lines) of known real structures in the Argo-SIM slide were employed as standard samples for 2D-SIM imaging (**Fig. 2g-t**). Since structure of the samples was known (rings and lines), raw data with high modulation and high SNR were collected with the exposure time of 50 ms, and SR images with minimal artifacts were reconstructed thereafter. The residual noise artifacts in the SR image were eliminated by setting thresholds, and clean SR images were obtained as the ground-truth models (**Fig. 2g,n**). Error maps and corresponding root mean square error (r.m.s.e) values, between the SR image and ground-truth image, were displayed to evaluate the fidelity of reconstruction algorithm. Further, Structure Similarity Index (SSIM) was used to quantitatively evaluate the fidelity of SR images, defined as

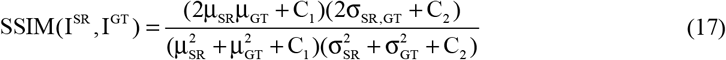

where μ_SR_ and μ_GT_ are the mean values of images I^SR^ and I^GT^ respectively, σ_SR_ and σ_GT_ are the standard deviations of I^SR^ and I^GT^ respectively, and σ_SR, GT_ is the cross-variance between images I^SR^ and I^GT^; C_1_ and C_2_ were used to avoid division by a small denominator, and set as C_1_ = 0.05 and C_2_ = 0.05.

For comparative analysis, SoftWoRx, fairSIM, and HiFi-SIM were used to reconstruct the ring sample data with 10-ms exposure time and line sample data with 5-ms exposure time. To quantitatively analyze the influence of Wiener constants, the spectrum-optimization Wiener constant w_1_ in HiFi-SIM was set to 0.9 and 1.2 respectively; the deconvolution Wiener constant of fairSIM and HiFi-SIM were set to 0.005, 0.01, 0.05, 0.1, 0.2, 0.3, 0.4, and 0.5, respectively. SoftWoRx only set a Wiener constant of 0.005 for reconstruction. Ten different regions of interest (ROI), containing ring or line structures, from the SR images were selected to calculate the SSIM values between ROI images and corresponding ground-truth images (**Fig. 2t**).

## Software availability

The version of HiFi-SIM used in this paper is available as **Supplementary Software**.

## Data availability

Data in support of the findings of this study are available from the corresponding authors upon reasonable request.

